# Inferring interactions from microbiome data

**DOI:** 10.1101/2023.03.30.534939

**Authors:** Román Zapién-Campos, Florence Bansept, Arne Traulsen

## Abstract

Parameter inference of high-dimensional data is challenging and microbiome time series data is no exception. Methods aimed at predicting from point estimates exist, but often even fail to recover the true parameters from simulated data. Computational methods to robustly infer and quantify the uncertainty in model parameters are needed. Here, we propose a computational workflow addressing such challenges – allowing us to compare mechanistic models and identify the values and the certainty of inferred parameters. This approach allows us to infer which kind of interactions occur in the microbial community. In contrast to point-estimate inference, the distribution for the parameters, our outcome, reflects their uncertainty. To achieve this, we consider as many equations for the statistical moments of the microbiome as parameters. Our inference workflow, which builds upon a mechanistic foundation of microscopic processes, can take into account that commonly metagenomic datasets only provide information on relative abundances and hosts’ ensembles. With our framework, we move from qualitative prediction to quantifying the likelihood of certain interaction types in microbiomes.

---

Numerous studies have shown how important the microbiome is for their hosts, ranging from development to health [1, 2]. The promise of manipulating the microbiome relies on having understood the ecological and evolutionary processes operating on it [3]. Although metagenomics studies have widely characterized microbiome samples [4], their connection to mathematical models and eco-evolutionary theories lags behind. Part of the gap is explained by an intrinsic difficulty in analysing microbiome data [5], in particular the inverse problem of robustly inferring model parameters – and thus interactions between microbes – from data (Fig.1 A). Pioneering studies have achieved point (“best guess”) estimates that allow accurate predictions of the dynamics [6], but fall short in estimating the true parameters from simulations [5]. This apparent contradiction, rooted in the high dimensionality of the parameter space and incomplete nature of data, has been discussed by Cao et al. [5]. Others have suggested to use Bayesian inference – where probabilities are assigned to parameter values – to go beyond point-estimates [7]. Here, we present a computational workflow that starts from defining microscopic transition rates in a mathematical model – describing ecological and evolutionary events (such as birth, migration, mutation or speciation) – and works with macroscopic statistical moments of the microbiome composition. Our Bayesian inference workflow, which naturally bypasses known limitations of point-estimate inference [5], is sufficiently flexible to test mathematical models as diverse as microbiome samples while quantifying the parameter uncertainty stemming from data limitations (Fig.1 C) – including its extrinsic noise. Two classical ecological models are used to illustrate its application on datasets describing absolute or relative abundances of microbes – namely, logistic growth and Lotka-Volterra models. The inference workflow outlined here bridges a gap between microbiome data and theoretical modelling.

## Results

### Developing an inference workflow

We propose a parameter inference workflow grounded on a mechanistic description of the dynamics of absolute abundances in a microbiome (Fig.1 C). First, we write down microscopic transition rates *T* describing changes in the microbiome composition of one host, the vector **n**, to other compositions (**n**^′^), given the parameter set ***θ***. Note that these rates can be functions of time *t* as well,

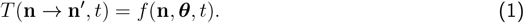

Now, instead of tracking the microbiome composition **n** in a single host, we can describe how the probability of observing a microbiome composition **n** in an ensemble of hosts, *P* (**n**, *t*), changes with time,

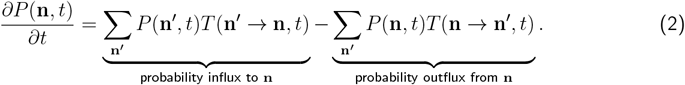

In this expression, called the master equation [8], the probability influx and outflux terms indicate an increase or decrease in the probability of composition **n**. This dynamics depends on the ecological and evolutionary processes contained in the microscopic transition rates.

Using the master equation, we can derive equations for the statistical moments of the microbiome composition in an ensemble of hosts – namely, the product of the master equation by a variable of interest (*g*_*k*_, where *k* is an identifying index) summed over all possible microbiome compositions **n**,

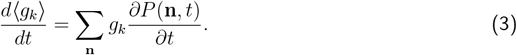

For example, computing the average abundance of microbial type *i* implies setting *g*_*k*_ = *n*_*i*_, whereas for the co-moment of microbial types *i* and *j, g*_*k*_ = *n*_*i*_*n*_*j*_. Equations like these describe the expected macroscopic dynamics of the microbiome. To obtain a system of equations of the moments that fully determines the parameters, we have to derive at least as many equations for them as parameters in the model. For example, in a Lotka-Volterra model with *S* microbial types, there are *S* growth rates and *S*^2^ intra- and inter-specific interactions, amounting to *S* + *S*^2^ parameters. Thus, we would need *S* +*S*^2^ equations. We can use the *S* equations for the first moments ⟨*n*_*k*_⟩, *S* for the second moments 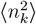, and *S*(*S* − 1) for the co-moments ⟨*n*_*k*_*n*_*l*_⟩ and covariances ⟨*n*_*k*_, *n*_*l*_⟩ (see Sup. Methods A). Note that each equation can depend on the vector of other moments, i.e., ⟨*g*_*k*_⟩ = *f* (⟨**g**⟩, ***θ***, *t*). A step by step derivation from microscopic rates up to second order moments for a logistic growth and the Lotka-Volterra models can be found in the Sup. Methods A. These models include conventional ecological events, such as growth, death, immigration, and direct and indirect interactions.

We now have the elements to infer the parameters ***θ*** from microbiome data. The idea of Approximate Bayesian Computation (ABC) is to identify feasible parameters values by comparing the data to model predictions [7]. Specifically, for a given set of parameters values ***θ***, a distance metric between the numerical solution of the equations for the moments, ⟨*g*_*k*_⟩, and the equivalent moments from data, 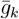, is estimated, e.g.,

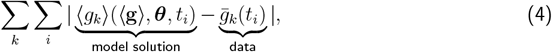

for the Euclidean distance. If this distance is smaller than a threshold *ε*, the set is considered valid. By testing sets of parameters sampled according to an expectation – prior distribution – and recording those below the threshold *ε*, a posterior distribution of the parameters reflecting the uncertainty of the inference can be obtained (Fig. 1 C). With a smaller threshold *ε* this posterior can become the new prior and the process can be iterated to improve it. This is called Approximate Bayesian Computation - Sequential Monte Carlo (ABC-SMC). We show how to choose prior distributions of the parameters in the Sup. Methods C.

**Figure 1.**
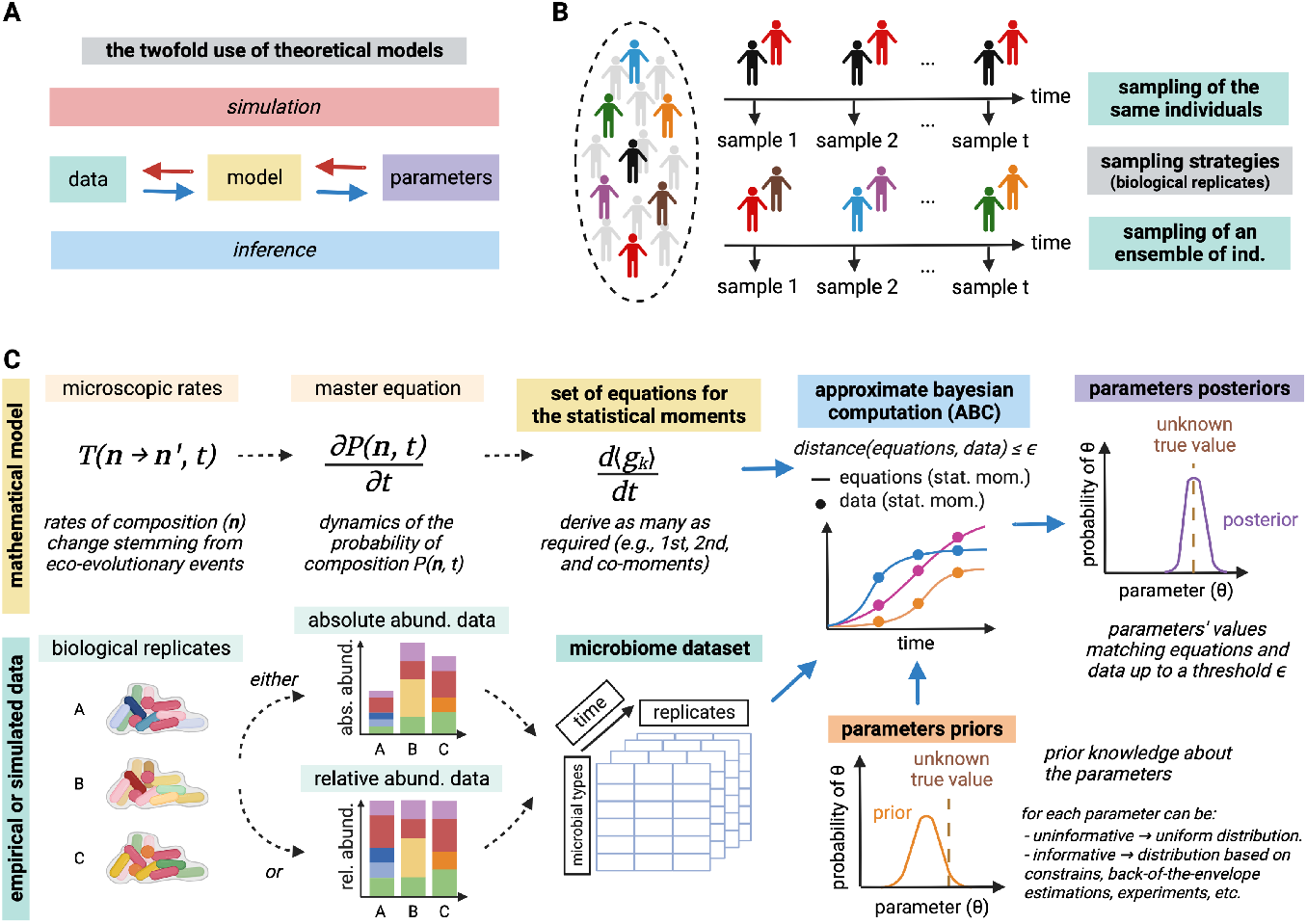
Parameter inference workflow proposed and microbiome data properties. (**A**) Mathematical models serve as a link between parameters and data. Either to simulate biological processes or to infer parameters from data. (**B**) Longitudinal sampling of the same hosts or an ensemble of them are used to obtain datasets. (**C**) Workflow from microscopic rates of a model and experimental data to inference of parameters values by Approximate Bayesian Computation (ABC). The microscopic rates describe possible eco-evolutionary events (such as birth, migration, mutation or speciation), leading to macroscopic patterns (statistical moments of the abundance). Datasets describe absolute abundances (counts) or relative abundances (frequencies) of microbes. To quantify the probability of parameter values given a dataset, prior knowledge about the parameters is updated to a posterior distribution based on the agreement of the model with the data.

### Properties of microbiome data

Given a microbiome dataset, all statistical moments 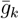 can be estimated from raw data. Concretely, it can be done by averaging the variable of interest *g*_*k*_, over all replicates in each specific time point (Fig. 1 C). For example, for *g*_*k*_ = *n*_*i*_, the replicates of *n*_*i*_ are simply summed over and divided by the number of replicates, while for *g*_*k*_ = *n*_*i*_*n*_*j*_, the products of *n*_*i*_ and *n*_*j*_ for each replicate are computed, then summed over and finally divided by the number of replicates.

Microbiome data is nowadays typically produced by metagenome sequencing. Conventionally, for technical reasons metagenomics only quantifies the *relative abundance* of each microbial type in a sample (Fig. 1 C) [9]. More recently, some studies have measured absolute numbers of culturable [10] and non-culturable microbes in samples [11]. We call these counts *absolute abundances*.

Our former equations only track moments of absolute abundance, ⟨*g*_*k*_⟩. As Gloor et al. [9] show, inferring parameters from relative abundance (*x*_*k*_) data using them would lead to spurious correlations (Fig. 2 A-B). To find equivalent expressions for the statistical moments of relative abundance, first, we propose *n*_Σ_ ≡ ∑_*j*_ *n*_*j*_ to be defined as a scaling factor and a dynamical equation for its first moment, ⟨*n*_Σ_⟩. Then, a transformation to moments of relative abundances, ⟨*γ*_*k*_⟩, is given by,

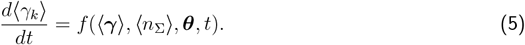

**Figure 2.**
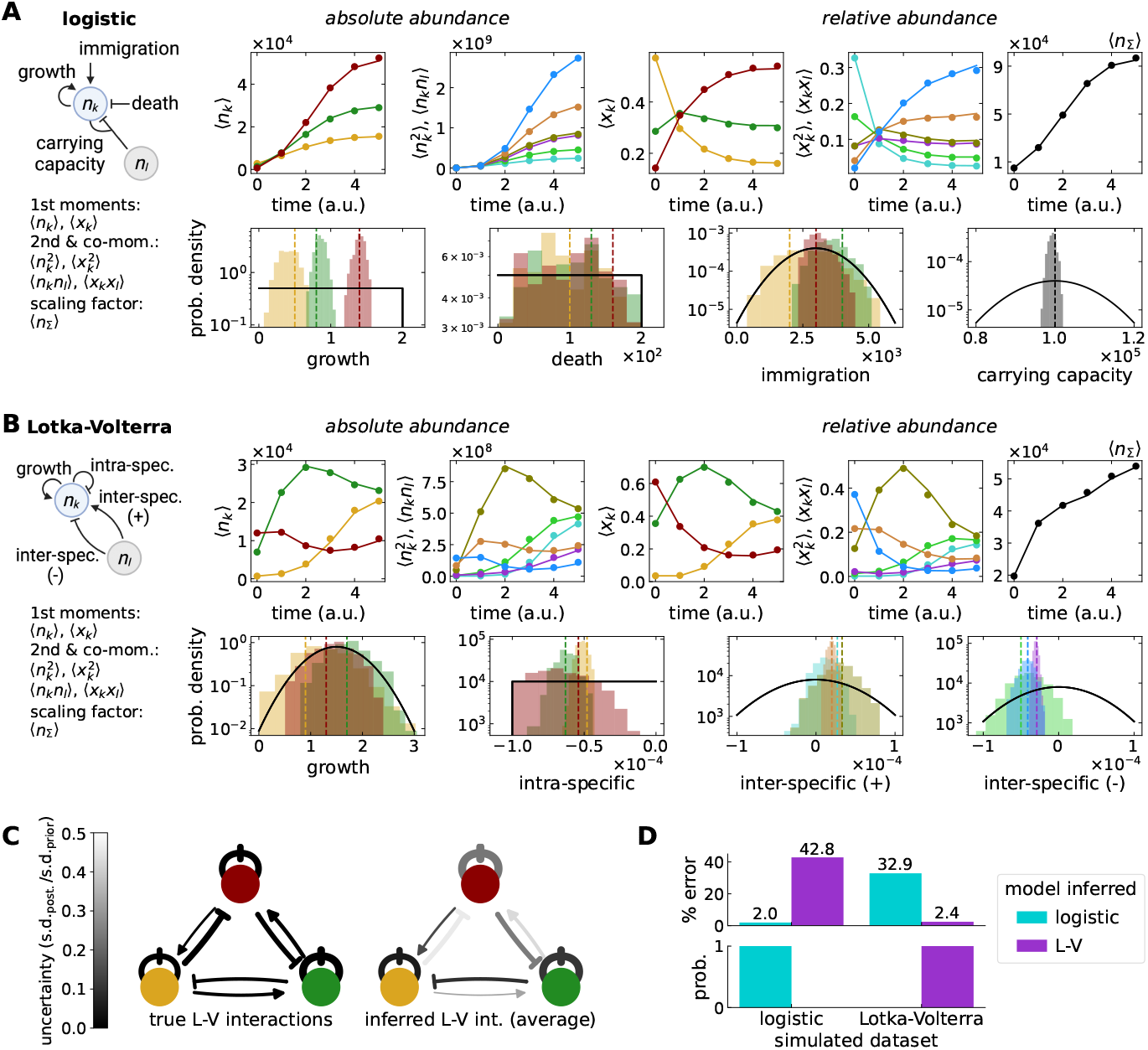
Inferring true parameters from simulated data. (**A-B**) Time series comparison between simulations (dots, derived from only 4 replicates) and equations for the statistical moments (lines) of absolute (*n*_*k*_) and relative abundance (*x*_*k*_) sharing parameters. Two models with three microbial types (*S* = 3) were tested, (**A**) logistic growth with immigration and death (3*S* + 1 = 9 parameters) and (**B**) Lotka-Volterra (*S* + *S*^2^ = 12 parameters). Inferred parameter posteriors from the relative abundance are compared to true parameters (dashed lines, Sup. Methods B) and priors (black distributions). All microbial types shared the same priors (Sup. Methods C). (**C**) The inferred interactions for the Lotka-Volterra model resembled the true interactions, qualitatively (arrow heads) and quantitatively (arrow thickness), with various certainties (grayscale, defined by the ratio of s.d. of posterior to prior). (**D**) For both datasets, the most probable model was identified correctly (a.u. = time units are determined by the rates, see Sup. Methods B).

Because relative abundance datasets lack information about the scaling factor, its initial condition, 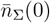, must be inferred as a free parameter, one parameter more than for absolute abundance data. Note that because ∑_*k*_ *x*_*k*_ = 1, the number of equations for the microbial types decreases but the number of parameters per type remains. A detailed derivation of transformations to relative abundance for a logistic growth and the Lotka-Volterra models is shown in the Sup. Methods A.

Apart from tracking the microbiome of the same hosts over time, as in animal gut studies, commonly, hosts sampled at different time points are pulled together to produce a single time series (Fig. 1 B). This is the case when hosts are sacrificed while sampling as in experimental studies of *D. melanogaster, C. elegans*, and *Hydra vulgaris* [10]. In contrast to deterministic models, our workflow can deal with hosts pulled together thanks to its account of the stochastic demography. More concretely – akin to the concept of biological replicates – if the parameter values and initial conditions are the same in each host sampled, we can account for their emerging demographic differences, i.e., differences in microbiome composition resulting from stochasticity.

### Inference from simulated and empirical data

We tested our inference workflow in two ways. Firstly, we recovered the correct parameters from simulated relative abundance data (Fig. 2). Our approach proved successful in cases with and without inter-specific interactions, namely, data simulated from logistic growth and Lotka-Volterra models. In both cases, our approach identified the correct model while in addition estimating their parameters values and certainty.

Secondly, we inferred parameters of an empirical time series of absolute abundance in a reduced mice microbiome – OMM^12^ [12] (Fig. 3). The posteriors suggested the growth rates of *Akkermansia muciniphila, Bacteroides caecimuris, Bifidobacterium longum*, and *Muribaculum intestinale* to be most certain, with average doubling times ranging from hours to days. Meanwhile, except from *B. caecimuris*, the average death and immigration rates were less certain, ranging from *≈* 4 · 10^5^ to· 10^6^ cells per day. Most of the certainties obtained from empirical data (Fig. 3) are smaller than those from simulations (Fig. 2), highlighting the limits of the model tested and inference from, noisy, empirical data. However, in each case we obtained a set of parameters – capturing interactions between microbes – with some level of certainty. To exemplify the utility of our outcome, here, our results point to selection as the ecological driver of the OMM^12^ dynamics, despite a possible compatibility of this data with a neutral hypothesis once it has reached steady state [13]. In this case, neutrality would imply that the parameter posteriors overlap between microbial types, which is not the case (Fig. 3).

**Figure 3.**
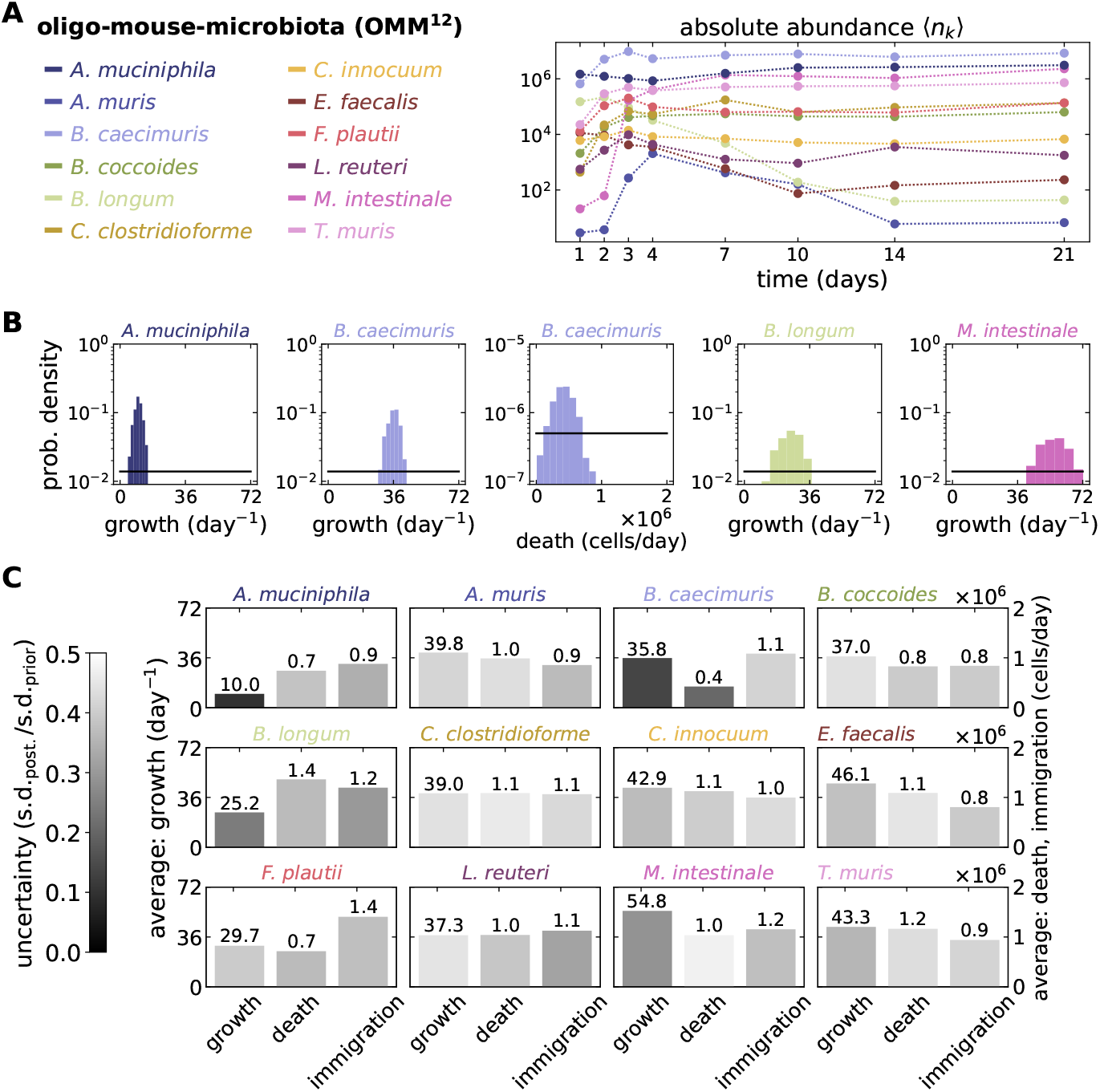
Inferring parameters from empirical data. The parameters of a logistic growth with immigration and death model were inferred from a mouse dataset. The Oligo-Mouse-Microbiota (OMM^12^) dataset [12] tracks a 12-species defined mice microbiome (*S* = 12), where the absolute abundances of the same individuals were sampled 11 times over 99 days. (**A**) We analysed the first 21 days where 4 replicates are available throughout and proposed a logistic growth model with parameters for growth, death and immigration. We illustrate 12 of the *S* + *S*^2^ = 156 moments used for the inference. (**B**) Of the 3*S* + 1 = 37 parameters inferred, we show only the posteriors of the five most certain ones (defined by the ratio of s.d. of posterior to prior). All microbial types shared the same uniform priors (black distributions, Sup. Methods C). (**C**) The parameters inferred for each species varied widely with various certainties. For the shared carrying capacity we found an average *N* ≈ 1.45 · 10^7^ bacterial cells, ±3.49 · 10^5^ cells and uncertainty of 0.0582. A system of 156 equations was solved (*S* = 12 1st moments and *S*^2^ = 144 2nd moments and co-moments).

## Discussion

The work presented here is motivated by the goal to understand how the microbes in a microbiome interact and the need to quantify the uncertainty of parameter estimation from microbiome data. Although inference methods for point estimation have been used [6], several issues limit their quantitative application, restricting them to recreate qualitative patterns of data [5]. A major issue is the indeterminacy of models with more parameters than equations [6, 5, 14]. We suggest a solution to this issue by deriving equations for the statistical moments of the microbiome composition – at least as many as parameters. In fact, our approach is driven by a mechanistic spirit, where microscopic rates must be written down first. As opposed to approaches where analytic solutions – or expensive stochastic simulations – are needed, here, a numerical solution is sufficient to quantify the distance to data [15]. This allows our workflow to handle very diverse models, where model comparison [15] in a Bayesian sense is possible.

The workflow is not limited by the properties of microbiome data [9]. As we have shown, analyzing datasets describing the relative abundance of microbial types – even if the total absolute abundance is dynamic [14] – is possible. Such is the nature of metagenomic sequencing data – the most common method to characterize microbiomes [5]. In addition, by tracking statistical moments of the microbiome, our approach naturally accounts for the diverse types of experimental samplings, such as those where ensembles of hosts are used to obtain a single time series. Concretely, compared to other methods, we track the demographic variation between hosts explicitly, and assign the remaining variation to external noise.

Although by design, our approach deals with longitudinal (time series) data, analyzing single time points (snapshot data) is possible. For example, if data is assumed to be at steady state, the inference method’s aim is to find parameters making the dynamical equations for the moments equal to zero. This does not mean that the moments are zero, but that their rate of change is. Nevertheless, as our results illustrate, given the various sources of uncertainty, time series data leads to better parameter inference, in particular, those time intervals of “high activity” where many changes occur [5]. As Cao et al. [5] proposed, several of these intervals could be analysed simultaneously to improve the inference.

Bayesian inference can suffer the curse-of-dimensionality in large and diverse systems [15]. By tracking numerous statistical moments readily solved numerically, we believe our approach combined with data of sufficiently high quality can overcome this to some extent – exploring the parameter space thoroughly in a reasonable time. We implemented an Approximate Bayesian Computation with Sequential Monte Carlo in our workflow using tools from the Python package pyABC [16] (Sup. Methods C), on which further optimizations could greatly improve its wider application [7]. As proof-of-principle, we applied our workflow to two simulated relative abundance datasets and recovered the true parameter values. We also applied it to a reduced microbiome in mice [12], where we estimated values and certainties of parameters describing logistic growth.

In summary, we presented a Bayesian inference workflow bridging microbiome data to theoretical modelling. By inferring datasets of microbial absolute and relative abundances, we showed its robustness – identifying likely interactions and certainty of parameters in simulated and empirical data. Because mechanistic rates serve as stepping stones of the workflow, similar microscopic models could replace the two classical ecological models that we illustrated – including experimentally informed models.

## Acknowledgements

We thank the *Theoretical Biology Department* in the MPI Plön and the *Collaborative Research Centre 1182: Origins and Functions of Metaorganisms* for the fruitful discussions. Finally, we thank the Max Planck Society and the CRC 1182 for the funding provided.

## Author contributions

RZC and AT developed the concept. RZC and FB worked on the methods. RZC wrote the first draft; all authors reviewed and approved the final version.

## Availability of data and software

The data generated and software used for the analyses are available in GitHub (https://github.com/romanzapien/microbiome-inference.git).

## Appendix

### A. Derivation of dynamical equations for the microbiome moments

To track the statistical moments of a model, e.g. average, variances, and co-variances, we have to account for the stochasticity of events. Thus, describing the probability of microbiome compositions is needed. The change in probability of each microbiome composition is described by the master equation,

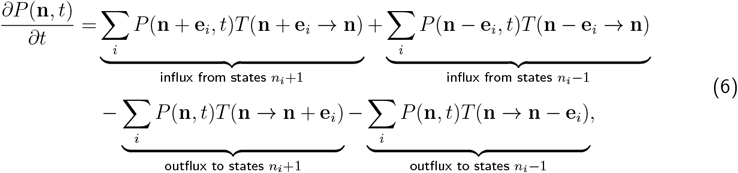

where **n** is the vector of absolute microbial abundances, and **e**_*i*_ is the amount of change, a vector with one in the *i-th* entry and zero otherwise.

Dynamical equations for the statistical moments can be obtained from the master equation by multiplication and subsequent summation. E.g., for the first moment ⟨*n*_*k*_⟩, equivalent to the average, we have

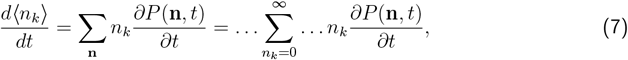

where for convenience, we make summations more explicit. For the second moment 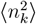, we have

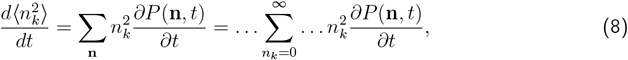

and for the co-moments ⟨*n*_*k*_*n*_*l*_⟩,

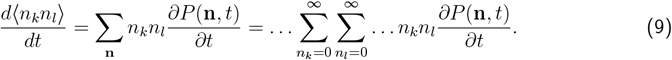

For models with a finite carrying capacity, the upper sum limit is changed to a finite number.

#### A.1. Logistic growth with immigration and death

Let us exemplify the former steps with a logistic growth model. Similarly to Allouche and Kadmon [17], let us define the microscopic transition rates for one microbial population *i*

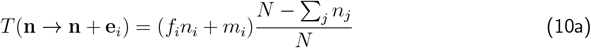

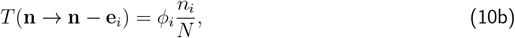

where *N* is the shared carrying capacity, *f*_*i*_ is the maximum growth rate, and *ϕ*_*i*_ and *m*_*i*_ are the death and immigration rates for each type *i*.

Now, we illustrate how to derive dynamical equations for the moments. Let us start with the first moment,

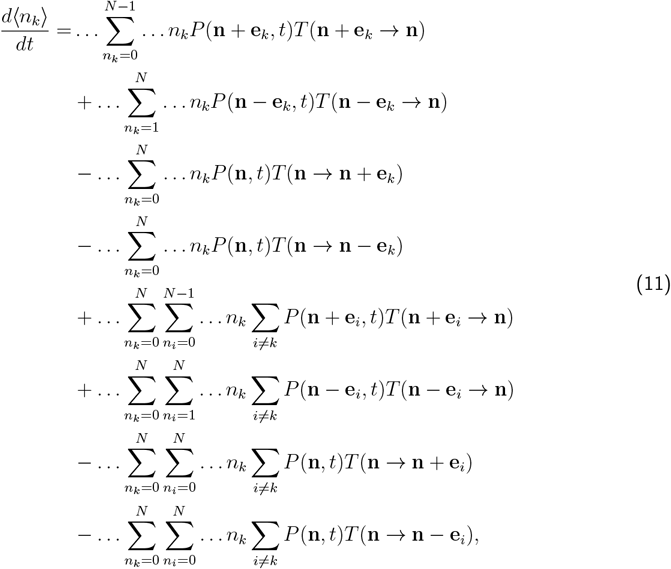

where the first four lines describe birth or death of a microbe of type *k* and the last four lines describe birth or death of a microbe of type *i* different from *k*. Note that by definition at the boundaries 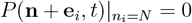 and 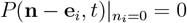, so their summation indices go up to *n*_*i*_ = *N* − 1, or start from *n*_*i*_ = 1, respectively.

After appropriate transformations of variables to only deal with *P* (**n**, *t*) and re-indexing, we obtain

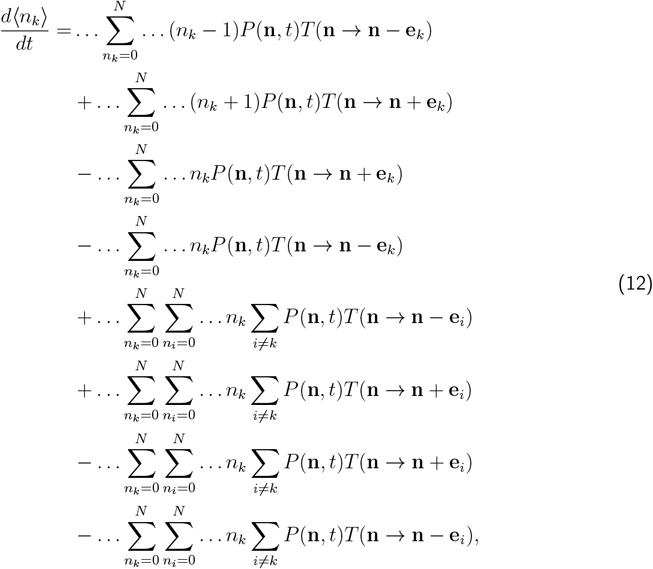

Note that the last four terms reduce to zero, and that at the boundaries 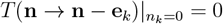 and 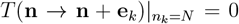, which allows including *n*_*k*_ = 0 and *n*_*k*_ = *N* in the summations. Simplifying, we find

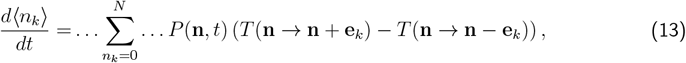

and substituting the transition rates *T* (**n** → **n** + **e**_*i*_) and *T* (**n** → **n** − **e**_*i*_) from Eqs. (10) leads to

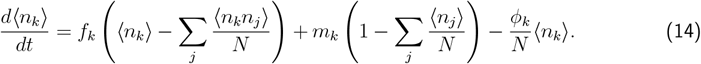

For other moments and models similar derivations can be done.

For the second moment, we find

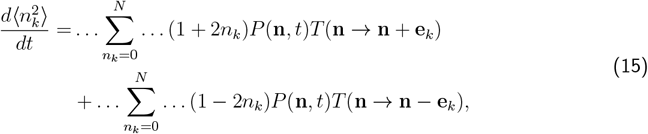

which after substituting *T* (**n** → **n** + **e**_*i*_) and *T* (**n** → **n** − **e**_*i*_) from Eqs. (10) reduces to

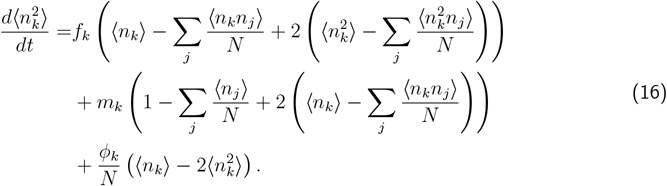

For the co-moments, we find

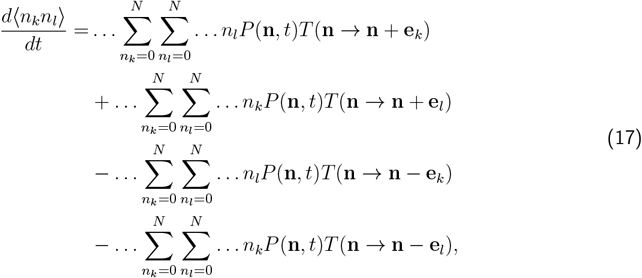

which after substituting *T* (**n** → **n** + **e**_*i*_) and *T* (**n** → **n** − **e**_*i*_) from Eqs. (10) leads to

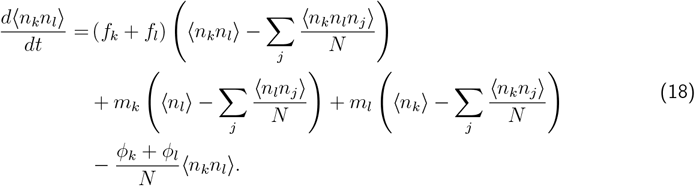

Because each equation depends on higher moments, e.g., *d*⟨*n*_*k*_*n*_*l*_⟩*/dt* depends on ⟨*n*_*k*_*n*_*l*_*n*_*j*_⟩, it is not possible to solve this system of equations without additional assumptions. However, one can find approximate expressions, where lower moments replace higher moments. For example, 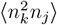 and ⟨*n*_*k*_*n*_*l*_*n*_*j*_⟩ are approximated as functions of the lower moments: 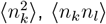, ⟨*n*_*k*_*n*_*l*_⟩, and ⟨*n*_*j*_⟩. Concretely, we can approximate e.g. ⟨*n*_*k*_*n*_*l*_*n*_*j*_⟩ *≈* ⟨*n*_*k*_*n*_*l*_⟩ ⟨*n*_*j*_⟩. This technique, called moment closure approximation, leads to a closed system of ODEs and we use it in our approach. Kuehn [18] makes a thorough review on this technique.

#### A.2. Lotka-Volterra

Now, for a model with intra- and inter-specific interactions, let us define the transition rates,

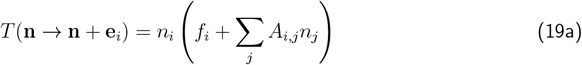

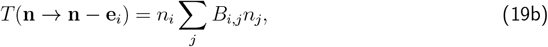

where *A* and *B* are positively defined matrices containing the interactions, satisfying *A*_*i,j*_ = 0 if *B*_*i,j*_ *>* 0, and *B*_*i,j*_ = 0 if *A*_*i,j*_ *>* 0. Ecologically, while interactions in *A* promote growth, those in *B* lead to death. Finally, *f*_*i*_ is the intrinsic growth rate.

For the first moment, similarly to Eq. (13), we have

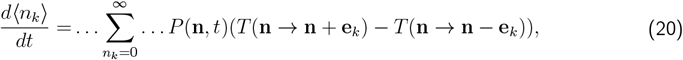

which after substituting *T* (**n** → **n** + **e**_*i*_) and *T* (**n** → **n** − **e**_*i*_) from Eqs. (19),

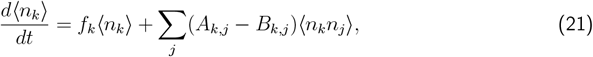

which takes the form of the conventional, determininistic Lotka-Volterra equations for the abundance with growth rate *f*_*k*_ and interaction matrix *A*_*k,j*_ − *B*_*k,j*_.

For the second moment, similarly to Eq. (15)

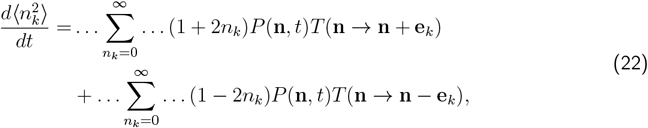

which after substituting *T* (**n** → **n** + **e**_*i*_) and *T* (**n** → **n** − **e**_*i*_) from Eqs. (19) leads to

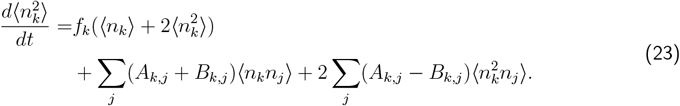

For the co-moments, similarly to Eq. (17), we derive

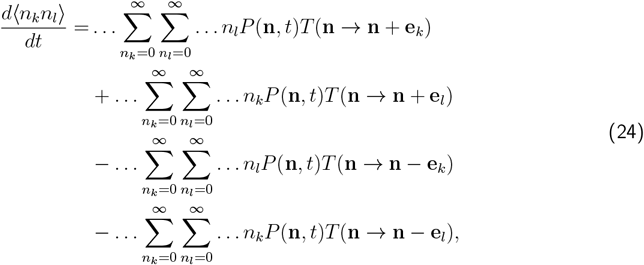

which after substituting *T* (**n** → **n** + **e**_*i*_) and *T* (**n** → **n** − **e**_*i*_) from Eqs. (19) reduces to

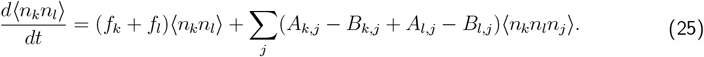

As previously, a moment closure approximation is required to solve the system of equations.

#### A.3. From absolute to relative abundance

The former equations account for the change of absolute abundances. To focus on relative abundance data, we define the relative abundance as

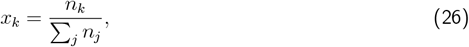

and

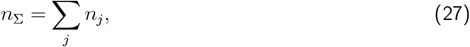

to serve as a scaling factor. Thus,

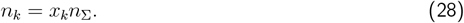

Let us find the transformation to relative abundances for the first moment. Using the definition of the covariance ⟨*x*_*k*_, *n*_Σ_⟩ = ⟨*x*_*k*_*n*_Σ_⟩ *−*⟨*x*_*k*_⟩ ⟨*n*_Σ_⟩, such that ⟨*x*_*k*_*n*_Σ_⟩ = ⟨*x*_*k*_⟩ ⟨*n*_Σ_⟩ + ⟨*x*_*k*_, *n*_Σ_⟩, we have

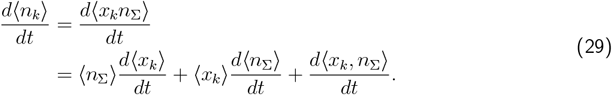

Rearranging, the transformation is given by

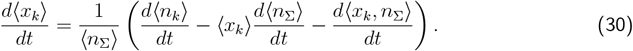

For second order moments, we use that 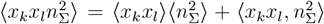 and approximate 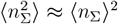. Then, using the chain rule

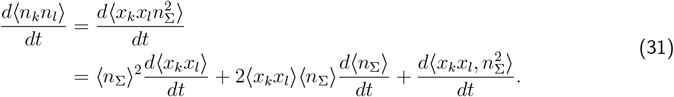

Rearranging, the transformations are given by

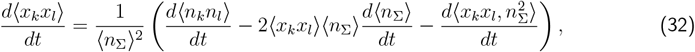

and

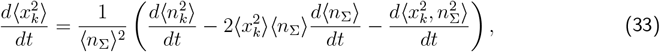

where the differential equation for ⟨*n*_Σ_⟩ is given by

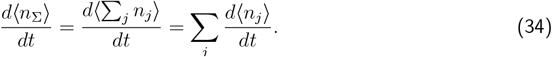

A close look at the dynamics of the covariances shows their contribution is negligible in large populations. To see this, let us write

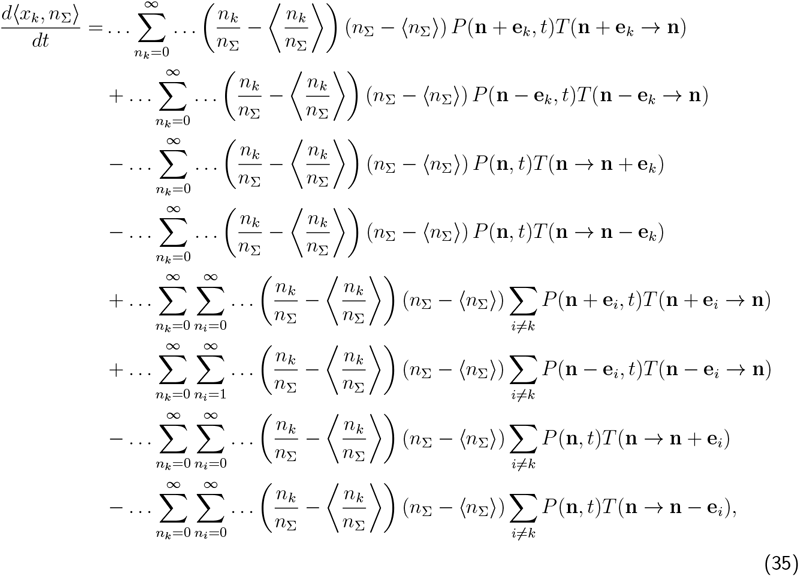

after the appropriate transformations of variable to only deal with *P* (**n**, *t*) and re-indexing, we find

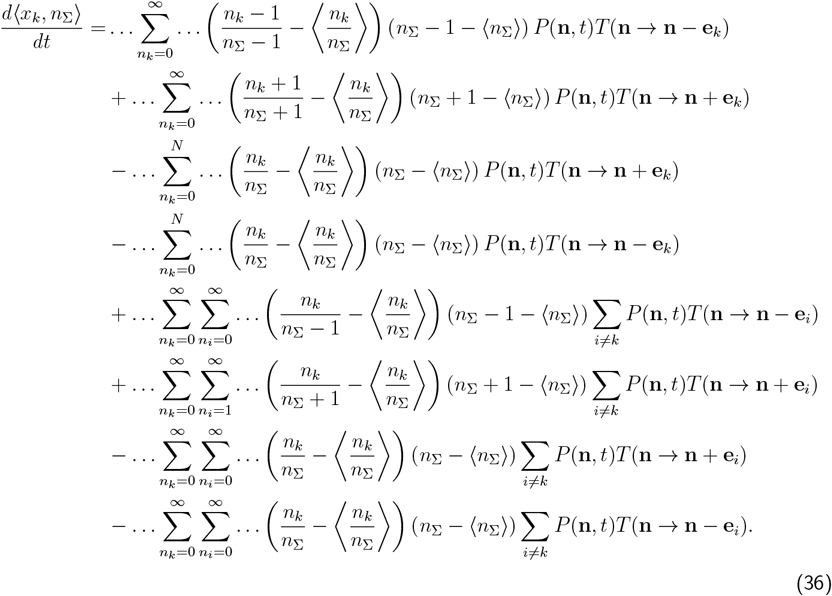

Note that if *n*_Σ_ ≫ 1, *n*_Σ_ *±* 1 *≈ n*_Σ_, then, if either *n*_*k*_ ≫ 1, such that *n*_*k*_ *±* 1 *≈ n*_*k*_ or 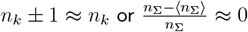, the terms from the previous equation simplify, leading to

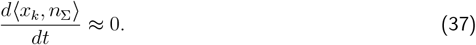

Similar arguments lead to conclude that,

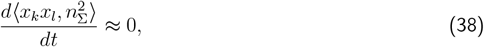

and

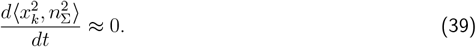

These approximations of the covariances are sensible in microbiomes, where *n*_Σ_, *n*_*k*_ ≫ 1 is often the case. Moreover, in the infinite population limit, covariances must be zero.

Putting all together, the change of the first moment of relative abundance in large populations is given by

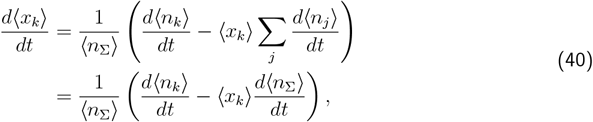

while for the second moments of relative abundance

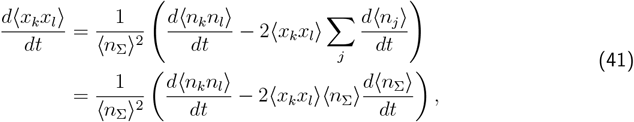

and

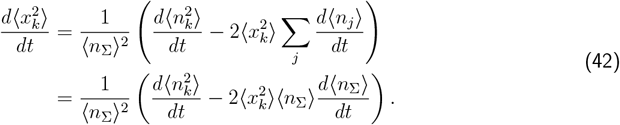

Finally, to solve these equations in terms of relative abundance, the change of variable *n*_*i*_ = *x*_*i*_*n*_Σ_ is needed all along. As Joseph et al. [14], we see that the second term of each equation serves as “correction factor” due to the fact that relative abundances must add up to one at all times.

### B. True parameters in simulations

**Table 1:**
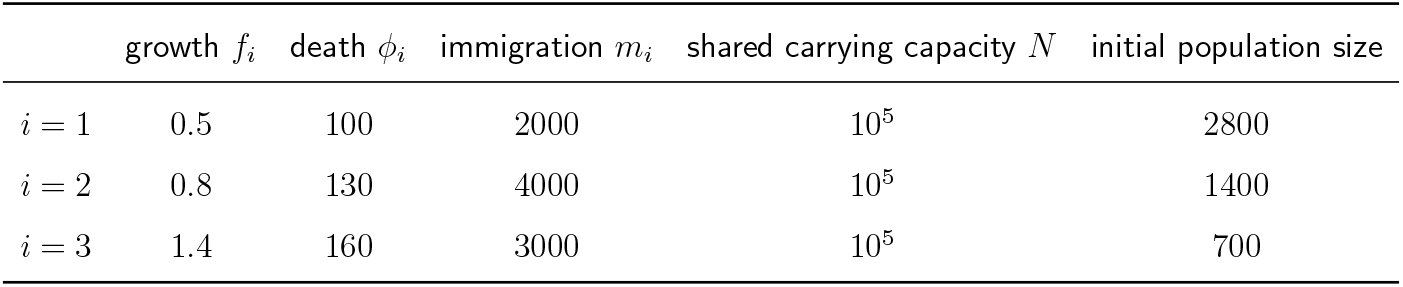
Parameters in simulated logistic growth with immigration and death (Fig. 2 A). The growth and death rates as well as the immigration parameters were only chosen for illustration, thus, time units are arbitrary.

**Table 2:**
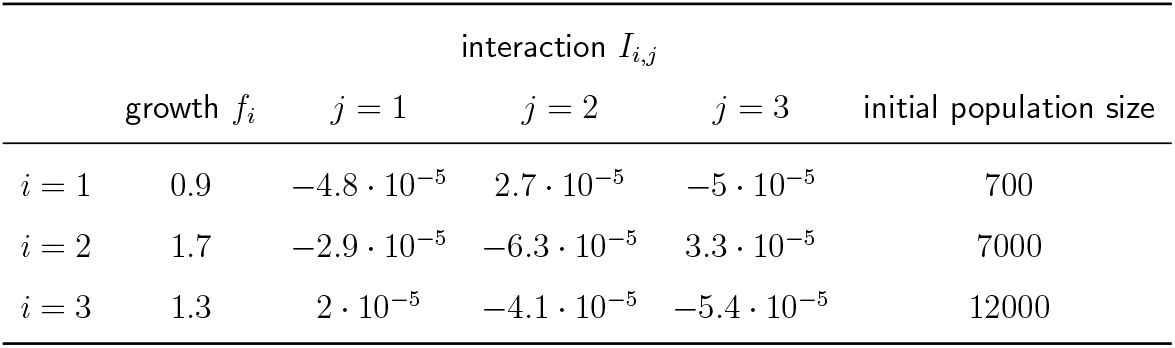
Parameters in simulated Lotka-Volterra (Fig. 2 B). The interaction parameters as well as the growth rates were only chosen for illustration, thus, time units are arbitrary.

| | | interaction $I_{i,j}$ | | | |
| --- | --- | --- | --- | --- | --- |
| | growth $f_i$ | $j = 1$ | $j = 2$ | $j = 3$ | initial population size |
| $i = 1$ | 0.9 | $-4.8 \cdot 10^{-5}$ | $2.7 \cdot 10^{-5}$ | $-5 \cdot 10^{-5}$ | 700 |
| $i = 2$ | 1.7 | $-2.9 \cdot 10^{-5}$ | $-6.3 \cdot 10^{-5}$ | $3.3 \cdot 10^{-5}$ | 7000 |
| $i = 3$ | 1.3 | $2 \cdot 10^{-5}$ | $-4.1 \cdot 10^{-5}$ | $-5.4 \cdot 10^{-5}$ | 12000 |

### C. TInference settings

**Table 3:** Priors for simulated logistic growth with immigration and death (Fig. 2 A). A combination of uninformative (uniform) and informative (normal) priors were used for illustration. These priors span a wide range of values to test the ability of the inference workflow to find the true parameters in simulations (Table 1). 𝒰 (*a, b*) indicates a uniform distribution in the range from *a* to *b*. 𝒩 (*a, b*) indicates a normal distribution with mean *a* and standard deviation *b*.

| parameter | units | prior |
| --- | --- | --- |
| growth $f_i$ | $\text{time}^{-1}$ | $\mathcal{U}(0, 2)$ |
| death $\phi_i$ | cells/time | $\mathcal{U}(0, 200)$ |
| immigration $m_i$ | cells/time | $\mathcal{N}(3000, 1000)$ |
| shared carrying capacity $N$ | cells | $\mathcal{N}(10^5, 10^4)$ |
| initial scaling factor $\bar{n}_\Sigma(0)$ | cells | $\mathcal{U}(4000, 6000)$ |

**Table 4:** Priors for simulated Lotka-Volterra (Fig. 2 B). A combination of uninformative (uniform) and informative (normal) priors were used for illustration. These priors span a wide range of values to test the ability of the inference workflow to find the true parameters in simulations (Table 2). 𝒰 (*a, b*) indicates a uniform distribution in the range from *a* to *b*. 𝒩 (*a, b*) indicates a normal distribution with mean *a* and standard deviation *b*.

| parameter | units | prior |
| --- | --- | --- |
| growth $f_i$ | $\text{time}^{-1}$ | $\mathcal{N}(1.5, 0.5)$ |
| intra-specific $I_{i,i}$ | $(\text{cells} \cdot \text{time})^{-1}$ | $\mathcal{U}(-10^{-4}, 0)$ |
| inter-specific $I_{i,j}$ | $(\text{cells} \cdot \text{time})^{-1}$ | $\mathcal{N}(0, 5 \cdot 10^{-5})$ |
| initial scaling factor $\bar{n}_\Sigma(0)$ | cells | $\mathcal{U}(1.5 \cdot 10^4, 2.5 \cdot 10^4)$ |

**Table 5:**
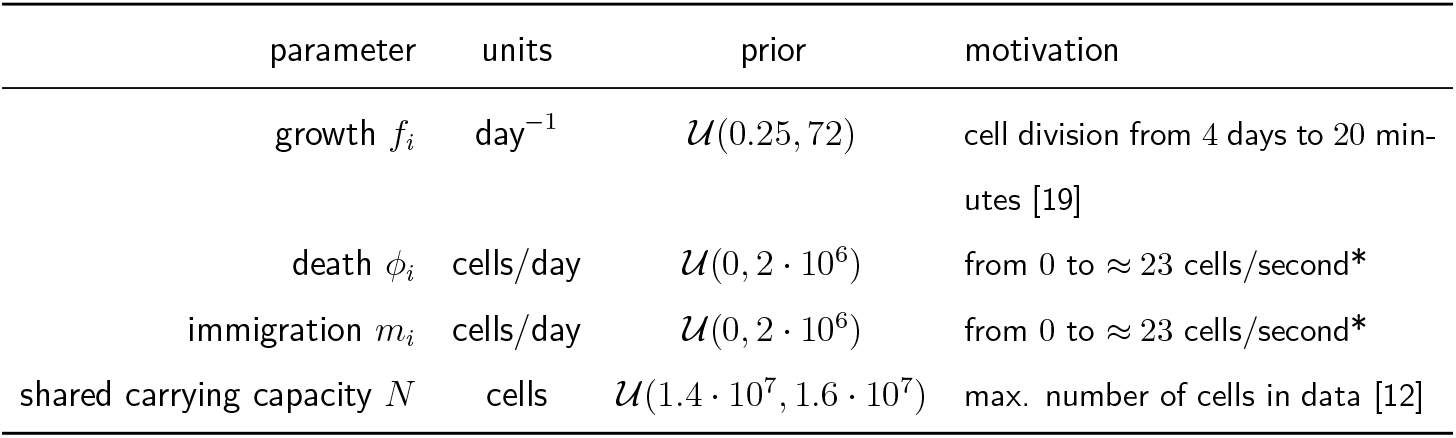
Priors for logistic growth with immigration and death in empirical mouse data (Fig. 3). Available evidence and back-of-the-envelope calculations (marked by *) were used to propose wide priors. 𝒰 (*a, b*) indicates a uniform distribution in the range from *a* to *b*. 𝒩 (*a, b*) indicates a normal distribution with mean *a* and standard deviation *b*.

| parameter | units | prior | motivation |
| --- | --- | --- | --- |
| growth $f_i$ | $\text{day}^{-1}$ | $\mathcal{U}(0.25, 72)$ | cell division from 4 days to 20 minutes [19] |
| death $\phi_i$ | cells/day | $\mathcal{U}(0, 2 \cdot 10^6)$ | from 0 to $\approx 23$ cells/second* |
| immigration $m_i$ | cells/day | $\mathcal{U}(0, 2 \cdot 10^6)$ | from 0 to $\approx 23$ cells/second* |
| shared carrying capacity $N$ | cells | $\mathcal{U}(1.4 \cdot 10^7, 1.6 \cdot 10^7)$ | max. number of cells in data [12] |

**Table 6:** Settings for Approximate Bayesian Computation - Sequential Monte Carlo (ABC-SMC) code. These settings were chosen to decrease the computing time, but still robustly minimize the distance between data and model. We used tools from the Python package pyABC [16], mainly *ABCSMC*. The maximum number of generations, mismatch threshold (*ε*) minimum, and minimum *ε* change between generations are all stopping criteria (marked by *). LSODA is a numerical solver capable of selectively adapting to the stiffness of a system of differential equations. *NA*: Not Applicable.

|  | simulations (Fig. 2) |  | mouse data (Fig. 3) |
| --- | --- | --- | --- |
|  | logistic | Lotka-Volterra | logistic |
| *numeric integration (numpy)* |  |  |  |
| method | LSODA |  | LSODA |
| *ABC-SMC (pyABC)* |  |  |  |
| calibration samples (to get first $\varepsilon$) | 2000 | | 400 |
| * max. number of generations | 15 |  | 10 |
| accepted samples per generation | AdaptivePopulationSize (CV = 0.4) |  | APS (CV = 0.25) |
| samples in first generation | 750 |  | 2000 |
| min. samples per generation | 500 |  | 500 |
| max. samples per generation | 1000 |  | 1000 |
| mismatch threshold ($\varepsilon$) update | QuantileEpsilon (best 10%) | | QE (10%) |
| strategy to sample parameters | MultivariateNormalTransition |  | MNT |
| * mismatch threshold ($\varepsilon$) min. | 0.046 | 0.071 | NA |
| * min. $\varepsilon$ change btw. generations | NA | | $10^{12}$ |

